# Using relative brain size as predictor variable: serious pitfalls and solutions

**DOI:** 10.1101/2022.03.15.484414

**Authors:** Simeon Q. Smeele

**Affiliations:** Cognitive & Cultural Ecology Research Group, Max Planck Institute of Animal Behavior, Radolfzell, Germany; Department of Human Behavior, Ecology and Culture, Max Planck Institute for Evolutionary Anthropology, Leipzig, Germany; Department of Biology, University of Konstanz, Konstanz, Germany

**Keywords:** Bayesian statistics, comparative analysis, multiple regression, relative brain size, structural equation model

## Abstract

1. There is a long-standing interest in the effect of relative brain size on other life history variables in a comparative context. Historically, residuals have been used to calculate these effects, but more recently it has been recognised that regression on residuals is not good practice. Instead, absolute brain size and body size are included in a multiple regression, with the idea that this controls for allometry.
2. I use a simple simulation to illustrate how a case in which brain size is a response variable differs from a case in which relative brain size is a predictor variable. I use the simulated data to test which modelling approach can estimate the underlying causal effects for each case.
3. The results show that a multiple regression model with both body size and another variable as predictor variable and brain size as response variable work well. However, if relative brain size is a predictor variable, a multiple regression fails to correctly estimate the effect of body size.
4. I propose the use of structural equation models to simultaneously estimate relative brain size and its effect on the third variable and discuss other potential methods.

## Introduction

Brain size is often theorised to be important in the life history of species and has been linked to many traits such as innovation rates (Lefebvre, Reader, and Sol 2004), sociality (Dunbar and Shultz 2007) and longevity (Minias and Podlaszczuk 2017). It is important to note that it is often the *relative* and not the *absolute* size of the brain that is of interest. To control for allometric scaling (larger species have larger brains), brain size is measured relative to body size. There exists a large diversity of relative brain size measures (Healy and Rowe 2007), but some decades ago the use of residual brain size was proposed and quickly became the most popular (Jerison 1973). This measure is the residual from a model that has log body size as predictor and log brain size as response variable. It is an intuitively attractive approach, since it measures whether the brain is smaller or larger than expected from allometry alone. Multiple challenges were recognised early on. Probably the most debated is the effect of phylogeny (Armstrong 1983). When comparing across multiple species nested at different levels of taxonomy, how does one estimate the ‘true’ slope of the model? Several techniques have been proposed to mitigate this, but the debate is ongoing (Font et al. 2019; Burger et al. 2019).

More recently another statistical caveat has been highlighted (for an overview see Freckleton (2002)). The use of residuals leads to biased estimates if a correlation exists between the predictor variables, as is often the case in the comparative literature. The proposed solution is to include body size in a multiple regression, which is very similar to the use of residual brain size, if - and only if - relative brain size is the response variable. This approach takes care of both the phylogenetic signal (since the model can include the phylogenetic variance-covariance matrix) and correct estimation of uncertainty. However, one major issue has been overlooked: if relative brain size is a predictor variable, including body size as a second predictor leads to incorrect inference. This becomes clear when looking at the causal structure of such a case. The best way to decide which variables to include in an analysis is the use of causal graphs (Glymour, Pearl, and Jewell 2016; Wright 1934). Laubach et al. (2021) presented a simple guide to draw Directed Acyclic Graphs (DAG), in which all variables are connected with arrows that indicate the direction of causality.

To point out the difference between the case with relative brain size as response vs predictor variable, I will present the underlying assumed causal structure visualised in Figure 1. To avoid discussion about directionality I have simply named the third variable *z*. This might be confusing, so I will give an example for both cases and explain the evolutionary causal structure. In the first case brain size is the response variable. The social brain hypothesis is possibly the most hotly debated version of this case. Here brain size is a function of body size and some measure of sociality (*z*). The idea is that social species experience evolutionary pressure to increase brain size to deal with complex social situations. First brought up by Jolly (1966), this hypothesis has since received much attention both with residual brain size and absolute brain size as response variable (for a review see Acedo-Carmona and Gomila (2016)). The second case, where relative brain size is a predictor variable, has received less attention, but several studies have attempted to show that relative brain size can cause longevity, with relatively larger brained species living longer (Street et al. 2017; Yu et al. 2018; DeCasien et al. 2018; Jiménez-Ortega et al. 2020). The understanding is that cognitive ability causes both increased brain size and the ability to survive longer. This might be unintuitive, since mechanistically cognition is caused by the brain, but in an evolutionary context it makes sense, since species that require more cognitive flexibility experience evolutionary pressure to increase brain size. Body size also causes an increase in longevity through several processes (e.g., reduced predation risk, lower metabolism).

**Figure 1:**
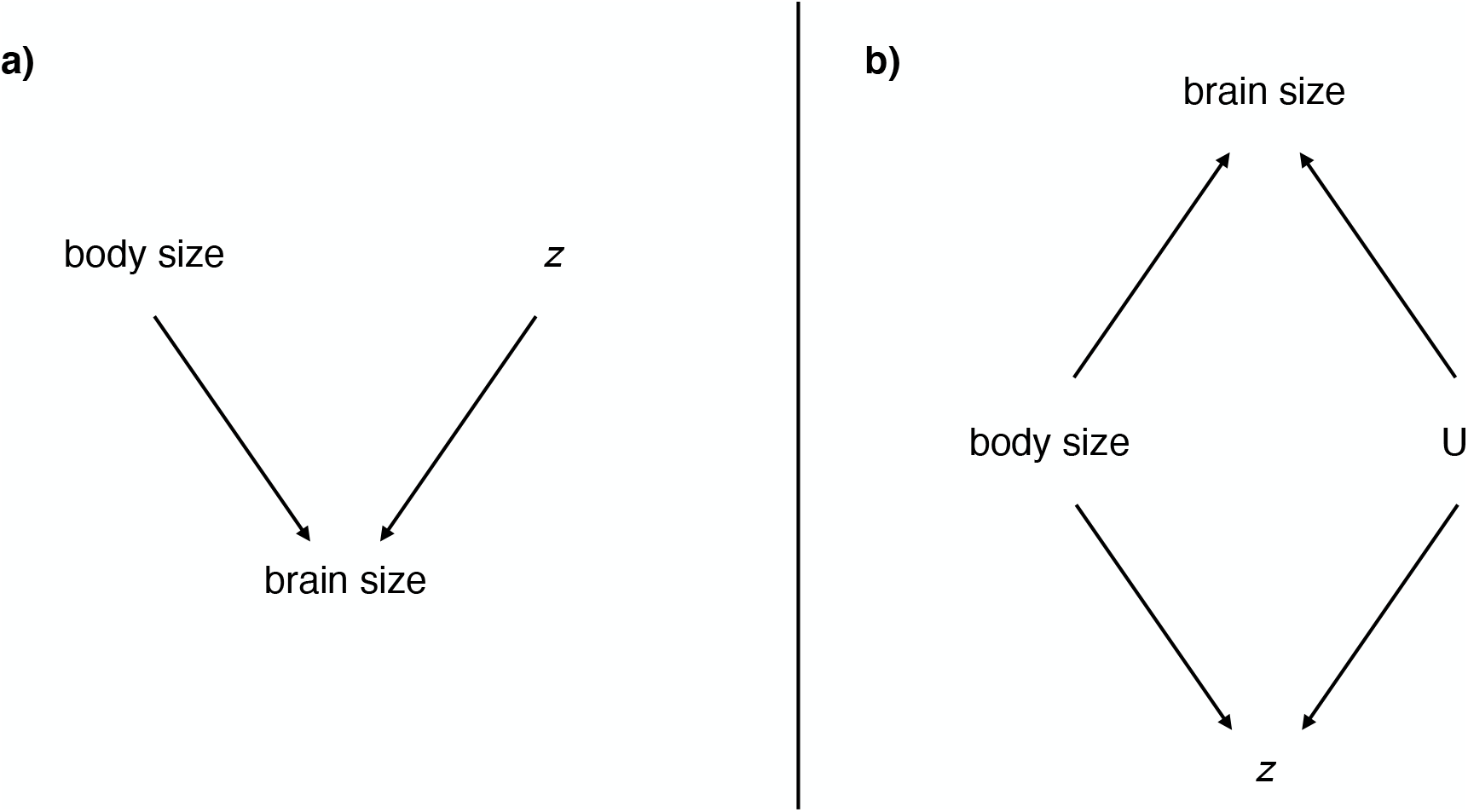
Directed Acyclic Graph of the two cases. Arrows indicate the direction of causality. z stands for the third variable of interest. a) Case I: brain size is the response variable. b) Case II: relative brain size is a proxy predictor variable. U stands for the unobserved cognitive ability.

In the first case both body size and *z* cause brain size (see Figure 1a). To decide how to best estimate the direct effect of *z* on brain size, one needs to make sure that all *back-door paths* are closed. In other words, all arrows that points towards *z* need to be considered. But since there are no such arrows in this case, there is no need to include additional variables. However, body size explains most of the variation in brain size. Therefore, it is still a good idea to include this variable to get a more precise estimate of the effect of *z* on brain size. Or in other words on relative brain size, since allometry is now accounted for.

The second case is causally more opaque (see Figure 1b). In general, the question of interest is how cognitive ability influences the third variable *z*. But since cognitive ability cannot be measured reliably across species (therefore denoted by a U for unobserved), relative brain size is used as a proxy. This is where it becomes tricky. The assumption is that it is the *extra bit of brain* that is caused by the need for cognitive ability. And it is therefore *relative* brain size that needs to be included as proxy. The variable we initially measured is *absolute* brain size, and is a collider in this DAG. A collider is a variable that is caused by two or more other variables. When including a collider in the analysis it *opens up* a back-door path, in our case through body size. This is problematic, because the estimated effect of brain size on *z* now also contains some of the effect of body size on *z*. Therefore, the estimated effect of body size on *z* will be biased. There is no way to correct for this using the variables in the DAG in a multiple regression. To include both relative brain size and body size one needs a system that contains regressions with brain size as response (of body size) and as predictor (of the third variable).

A structural equation model is such a system. It contains regressions for each variable and allows brain size to be response and predictor variable simultaneously (Bowen and Guo 2011). When fitted using a Bayesian approach, information flows in both directions, since the likelihood is computed for the whole system at each step. The aim of this paper is to show the estimation bias in multiple linear regressions using a simulation and propose a simple Bayesian structural equation model as a solution that can be easily adapted to most comparative studies. I also provide a version of the structural equation model with *ulam* from the *rethinking* package as front-end (McElreath 2020) so that models can be adapted within the R environment.

## Methods

To show the difference between a case where relative brain size is the *response* variable and where it is a *predictor* variable I simulated two simple cases with three variables (all code is publicly available at https://github.com/simeonqs/Using_relative_brain_size_as_predictor_variable-_serious_pitfalls_and_solutions). To simplify the simulation, I have not attempted to simulate realistic values for body size and relative brain size, but just used values with mean = 0 and standard deviation = 1. This allows me to draw general conclusions that are not sensitive to the scale of the variables. It should also be noted that body and brain size are normally measured on the natural scale, but log-transformed before analysis. The assumption is that causal effects are linear on the log-log scale. I have therefore simulated both body and brain size to be on the logarithmic scale.

I simulated 20 datasets per case and present parameter estimates for all datasets. Frequentist linear models were fitted with the *lm* function from R (R Core Team 2021). Bayesian models were fitted using the *cmdstanr* package (Gabry and Češnovar 2021), which runs the No U-turn Sampler in Stan (Gelman, Lee, and Guo 2015) with four chains and default sampler settings (1000 warmup and 1000 sampling iterations). Rhat, divergent transitions and effective sample size were monitored by the package and if issues arose these were reported in the results.

To test if sample size had an effect on which model performed best, additional simulations were run and analysed for 20 and 1000 species (see Appendix). To test if more or less informative priors had an effect on which model performed best, and how often the best model included the true effect in the credible interval, additional simulations were run with informative and vague priors (see Appendix).

### Case I: relative brain size as response variable

In the first case absolute brain size is caused by body size and *z*. The interest of the study is to what extent *z* causes additional variation in brain size (when the effect of body size is accounted for). In other words, to what extent *z* correlates with relative brain size. I simulated 20 data sets with 100 species with the following structure:

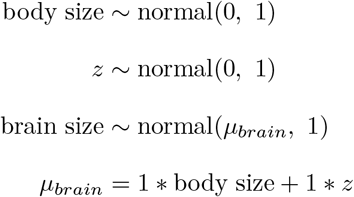

I analysed the resulting data with a frequentist linear model and with the Bayesian equivalent. Then I plotted the estimated coefficient of body size and *z* to show how well parameters were retrieved. Additionally, I reported the bias (difference between estimate and simulated value) and coverage (proportion of times the true value was within 2SE distance from the frequentist estimates and 2SD from the Bayesian estimates).

To test how models performed with an additional path from body size to *z*, I ran an additional simulation with this causal structure (see Appendix).

### Case II: relative brain size as predictor variable

In the second case both body size and relative brain size are a predictor of *z*. For simplicity I removed the unobserved variable from the simulation, and instead included a direct effect of relative brain size on z and absolute brain size. In this way absolute brain size is the sum of the brain tissue needed for enervating the body and the additional brain tissue, which is ultimately caused by other variables. Such a situation is still realistic when considering developmental time as a function of body size and relative brain size. The interest of the study is therefore to what extent relative brain size causes *z*. I simulated 20 data sets with 100 species with the following structure:

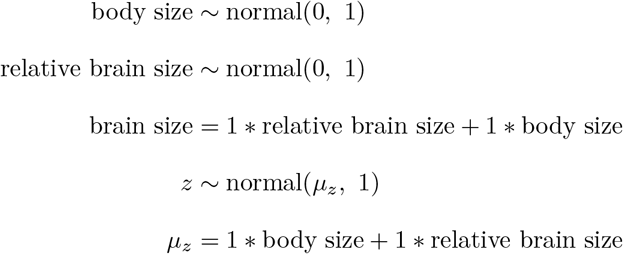

I analysed the resulting data with both a frequentist and Bayesian linear model where brain size and body size were included as predictor variables. Additionally, I analysed the data with a Bayesian structural equation model that included sub-models for all causal paths:

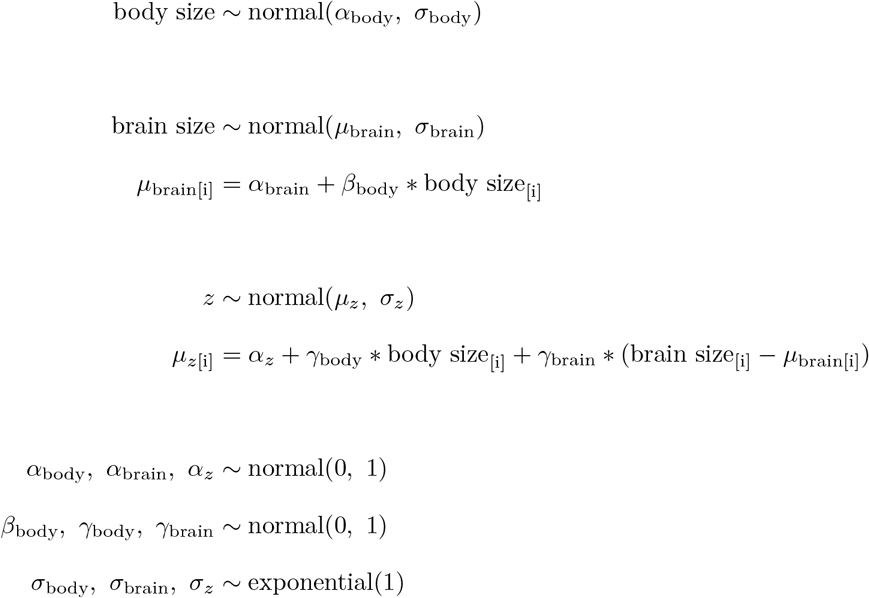

The model includes a regression for each variable. Body size is not a function of any variable. Brain size is a function of body size. *z* is a function of body size and relative brain size (where relative brain size is the difference between the actual and predicted brain size). Relative brain size in this model is very similar to residual brain size, but since it is computed at each iteration information flows in both directions and measurement error is correctly estimated. The last three lines of the model are the priors for all parameters. Note that the priors for the slopes (*β* and *γ*) are set to normal(0, 1), which regularises them slightly and centre them around no effect of the predictors. For empirical studies theory might provide more informative priors, which would further increase the accuracy of the model.

To test how models performed in case of unequal magnitude of the effect of body size and brain size on *z* (case II), additional simulations were run (see Appendix) with either strong body size effect (*β*_body_ = 2, *β*_brain_ = 0.5) or strong brain size effect (*β*_body_ = 0.5, *β*_brain_ = 2).

To visualise the results, I plotted all parameter estimates of the main model (for *z*) and reported the bias (difference between estimate and simulated value) and coverage (proportion of times the true value was within 2SE distance from the frequentist estimates and 2SD from the Bayesian estimates).

## Results

For Case I, where relative brain size was the response variable, both models estimated the parameters very well (see Figure 2, Table 1), also with an additional path from body size to *z* (see Appendix, Figure 14). For Case II, where relative brain size was a predictor variable, the effect of brain size was estimated well by all models, but the effect of body size was only estimated correctly by the structural equation model (see Figure 3, Table 1). Both the frequentist and Bayesian linear model estimated the effect of body mass to be essentially 0.

**Figure 2:**
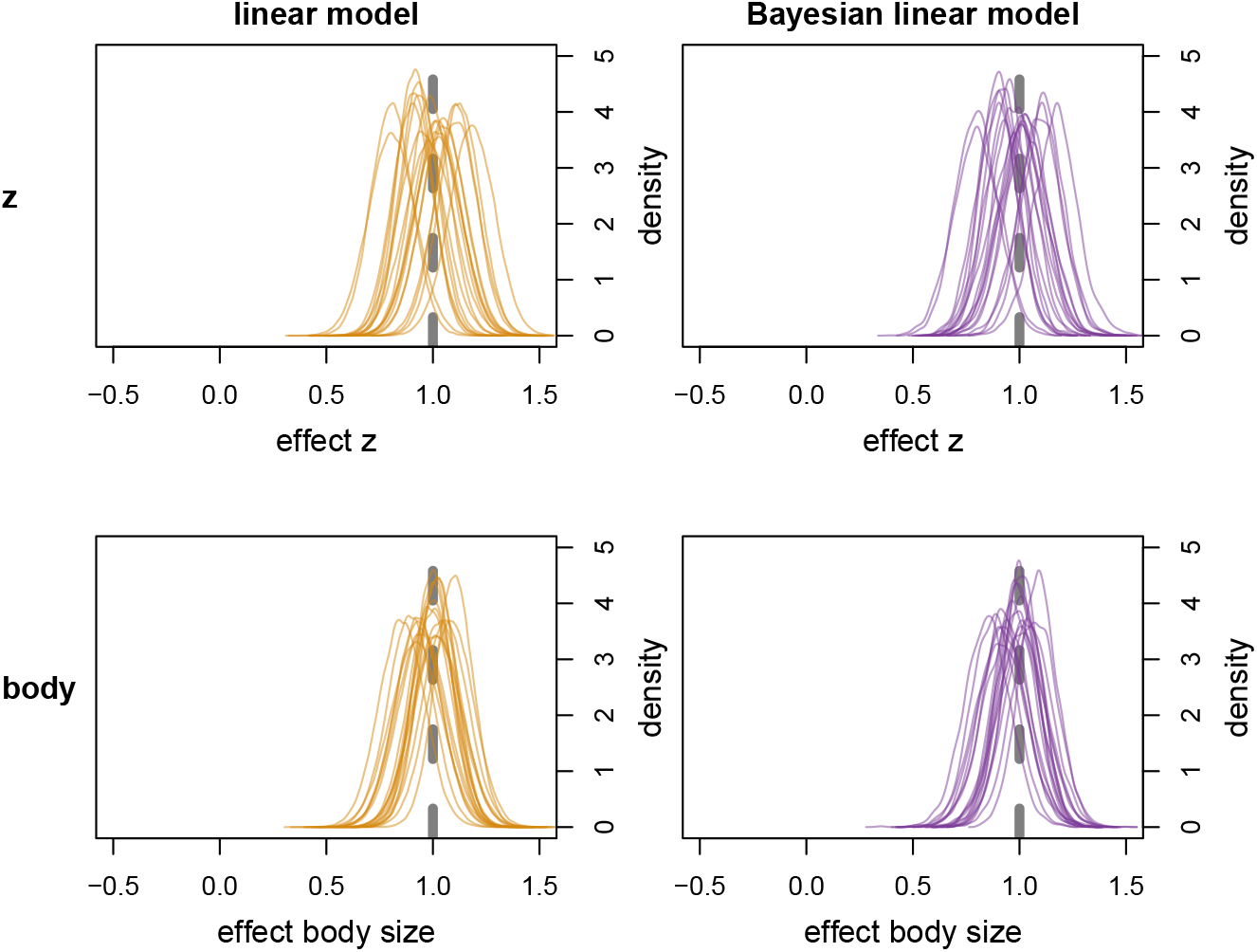
Parameter estimates from the linear model and Bayesian linear model with brain size as response variable. Dashed grey line is the true value. Orange density plots are normal distributions based on the mean and SE from the linear model. Purple density plots are the posterior distributions from the Bayesian model.

**Table 1:** Bias and coverage of all models. The mean and standard deviation of the bias is calculated from the distance between the point estimate and simulated value. Coverage is the proportion of times that the simulated value was withing the point estimate +− 2SE/SD. N is number of replicates.

| case | parameter | model | mean_bias | sd_bias | coverage | n |
| --- | --- | --- | --- | --- | --- | --- |
| response | body | linear | 0.01 | 0.06 | 1.00 | 20 |
| response | body | Bayesian linear | 0.02 | 0.06 | 1.00 | 20 |
| response | z | linear | 0.01 | 0.10 | 1.00 | 20 |
| response | z | Bayesian linear | 0.02 | 0.10 | 0.95 | 20 |
| predictor | body | linear | 1.03 | 0.16 | 0.00 | 20 |
| predictor | body | Bayesian linear | 1.02 | 0.16 | 0.00 | 20 |
| predictor | body | SEM | 0.08 | 0.16 | 0.90 | 20 |
| predictor | brain | linear | -0.02 | 0.10 | 0.95 | 20 |
| predictor | brain | Bayesian linear | -0.01 | 0.10 | 0.90 | 20 |
| predictor | brain | SEM | -0.01 | 0.10 | 0.90 | 20 |

**Figure 3:**
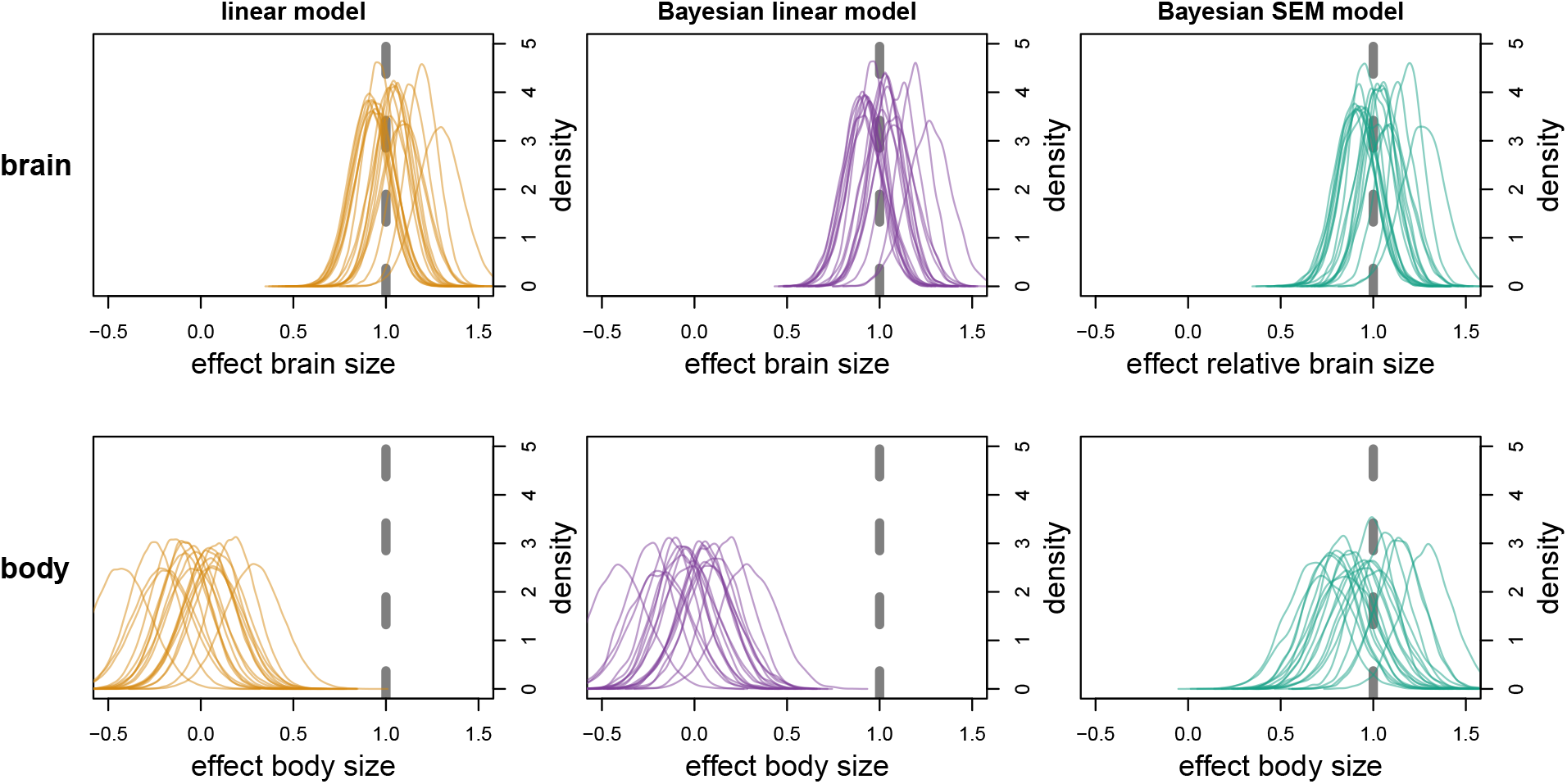
Parameter estimates from the linear model, Bayesian linear model and Bayesian strucutral equation model with relative brain size as predictor variable. Dashed grey line is the true value. Orange density plots are normal distributions based on the mean and SE from the linear model. Purple and green density plots are the posterior distributions from the Bayesian models.

Results were similar for simulations with smaller and larger sample sizes (see Appendix, Figure 4–7). Results were also similar for models with vague and informative prior (see Appendix, Figure 8–11). The informative priors constrained the Bayesian linear models estimate of the body size effect to positive values, but none of the posterior distributions included the true value of the body size effect. In other words, even if the model is run with the prior centred tightly around the correct value, it estimates the effect to be absent. Also when varying the magnitude of the effects of body size and brain size, the structural equation model was the only model to correctly estimate the effect of body size (see Appendix, Figure 12–13).

**Figure 4:**
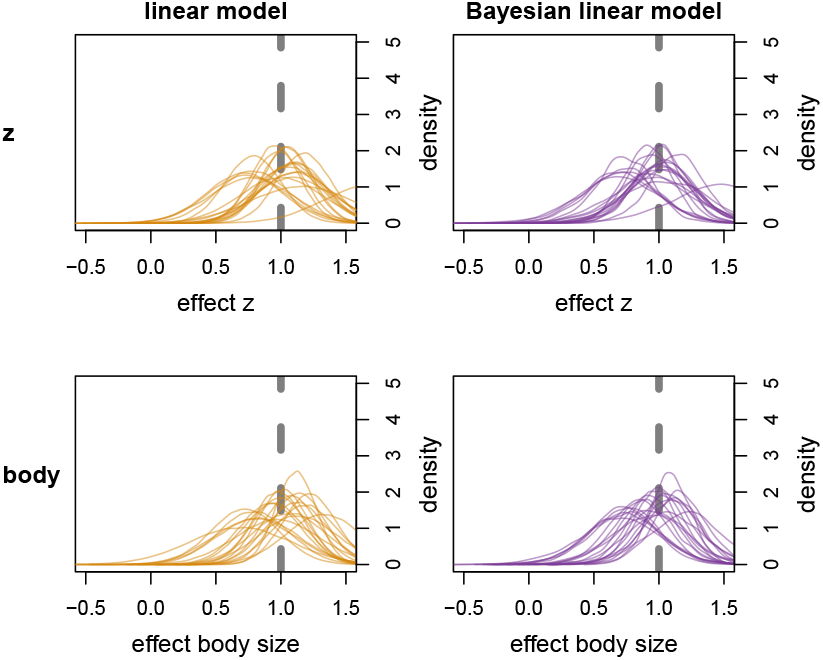
Model with sample size 20. Parameter estimates from the linear model and Bayesian linear model with brain size as response variable. Dashed grey line is the true value. Orange density plots are normal distributions based on the mean and SE from the linear model. Purple density plots are the posterior distributions from the Bayesian models.

**Figure 5:**
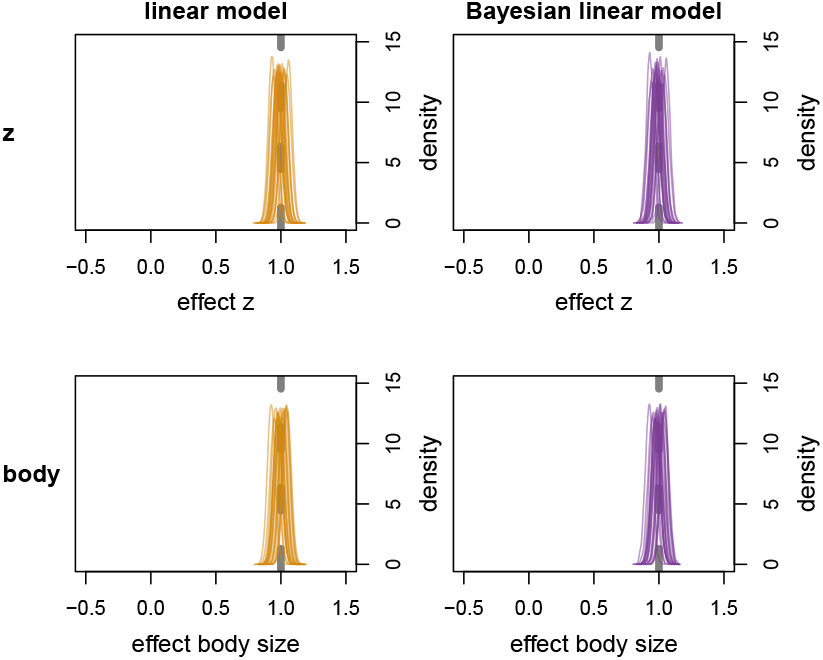
Model with sample size 1000. Parameter estimates from the linear model and Bayesian linear model with brain size as response variable. Dashed grey line is the true value. Orange density plots are normal distributions based on the mean and SE from the linear model. Purple and green density plots are the posterior distributions from the Bayesian models.

**Figure 6:**
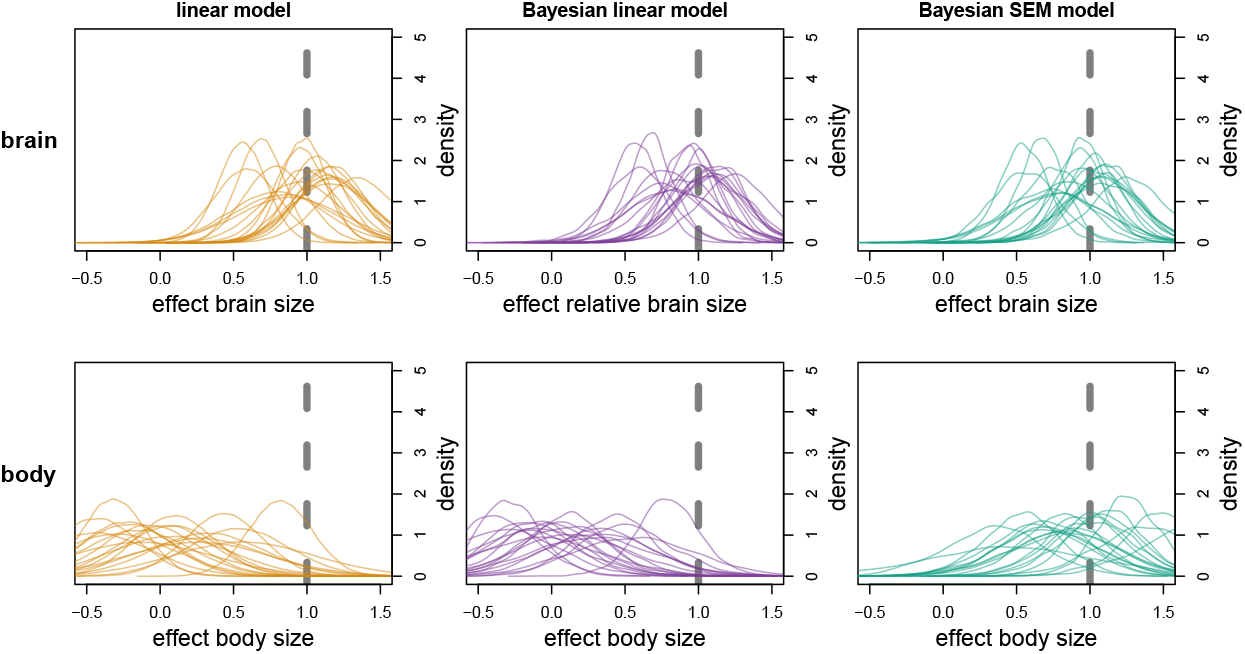
Model with sample size 20. Parameter estimates from the linear model, Bayesian linear model and Bayesian strucutral equation model with relative brain size as predictor variable. Dashed grey line is the true value. Orange density plots are normal distributions based on the mean and SE from the linear model. Purple density plots are the posterior distributions from the Bayesian models.

**Figure 7:**
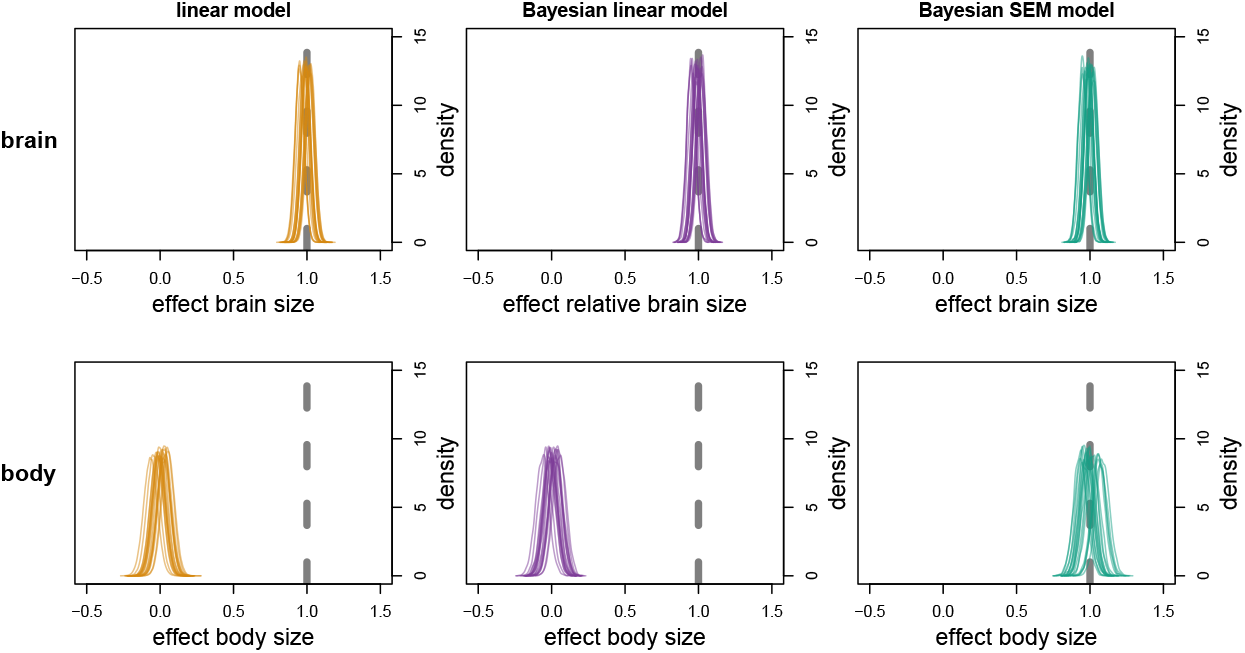
Model with sample size 1000. Parameter estimates from the linear model, Bayesian linear model and Bayesian strucutral equation model with relative brain size as predictor variable. Dashed grey line is the true value. Orange density plots are normal distributions based on the mean and SE from the linear model. Purple and green density plots are the posterior distributions from the Bayesian models.

**Figure 8:**
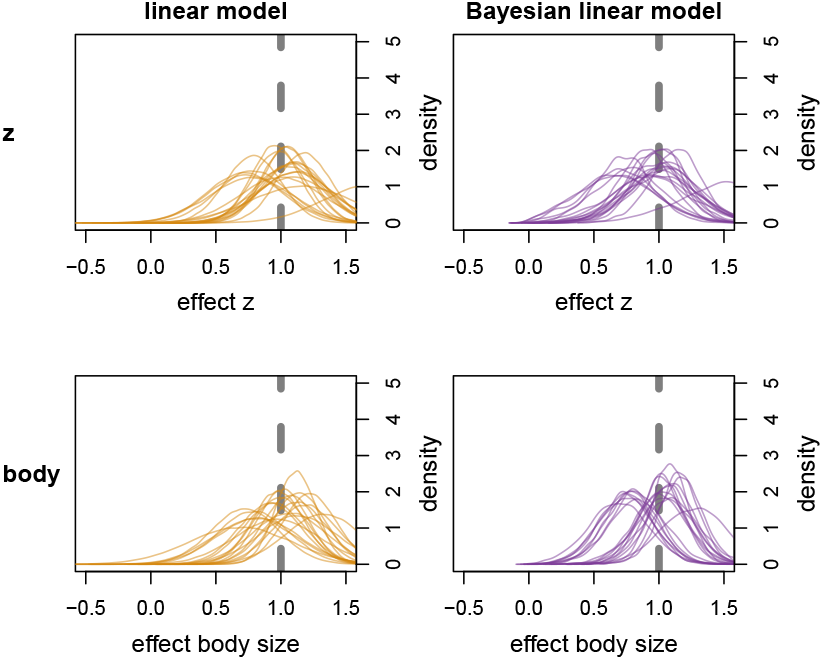
Model with informative priors. Parameter estimates from the linear model and Bayesian linear model with brain size as response variable. Dashed grey line is the true value. Orange density plots are normal distributions based on the mean and SE from the linear model. Purple density plots are the posterior distributions from the Bayesian models.

**Figure 9:**
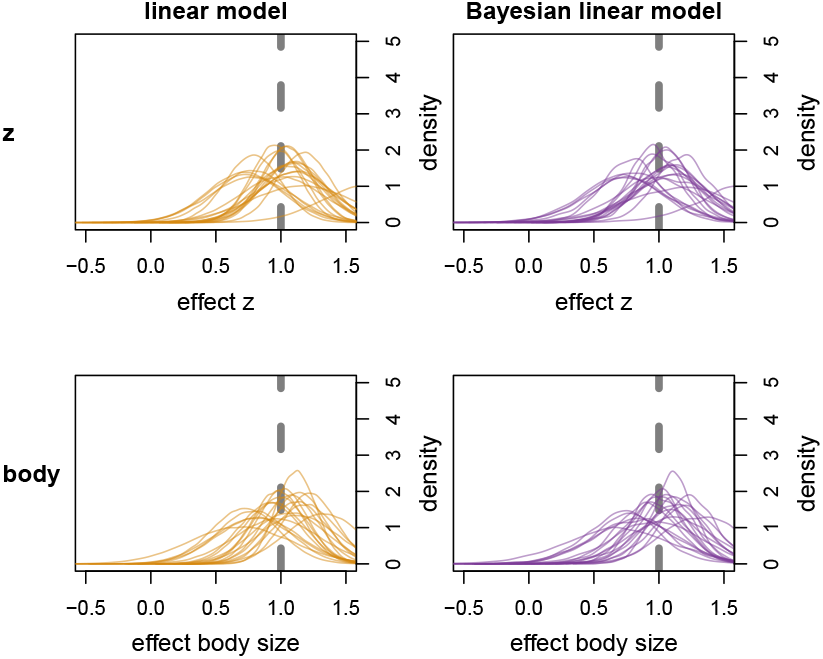
Model with vague priors. Parameter estimates from the linear model and Bayesian linear model with brain size as response variable. Dashed grey line is the true value. Orange density plots are normal distributions based on the mean and SE from the linear model. Purple and green density plots are the posterior distributions from the Bayesian models.

**Figure 10:**
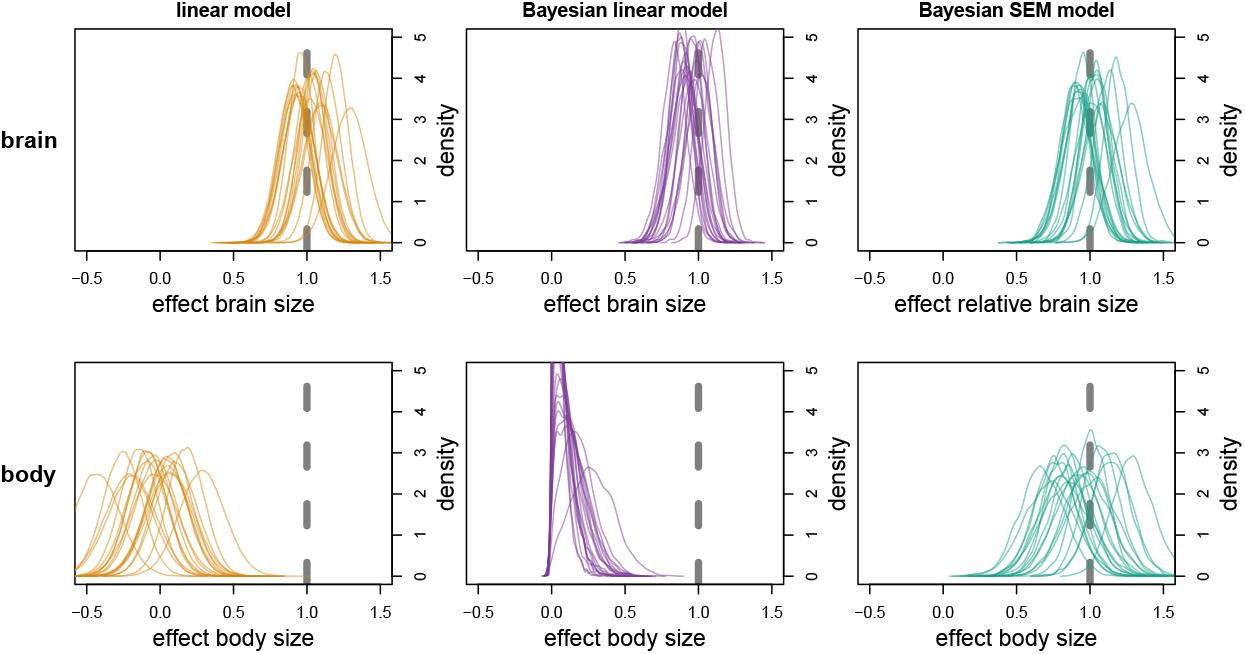
Model with informative priors. Parameter estimates from the linear model, Bayesian linear model and Bayesian strucutral equation model with relative brain size as predictor variable. Dashed grey line is the true value. Orange density plots are normal distributions based on the mean and SE from the linear model. Purple density plots are the posterior distributions from the Bayesian models.

**Figure 11:**
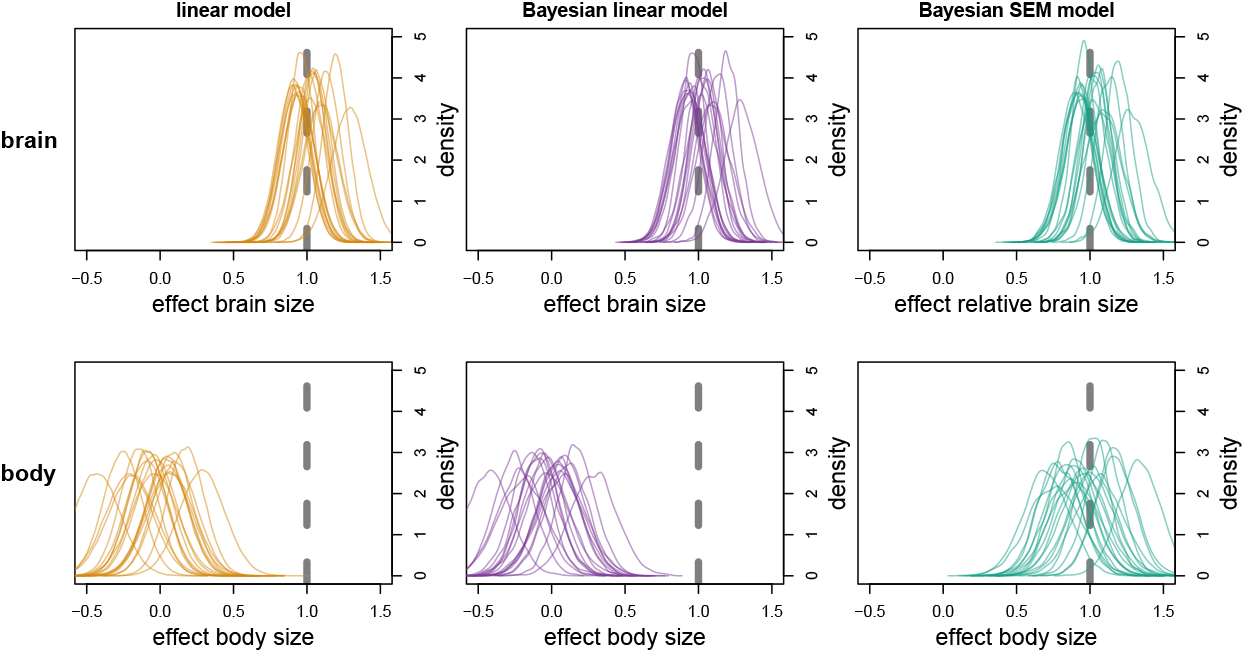
Model with vague priors. Parameter estimates from the linear model, Bayesian linear model and Bayesian strucutral equation model with relative brain size as predictor variable. Dashed grey line is the true value. Orange density plots are normal distributions based on the mean and SE from the linear model. Purple and green density plots are the posterior distributions from the Bayesian models.

**Figure 12:**
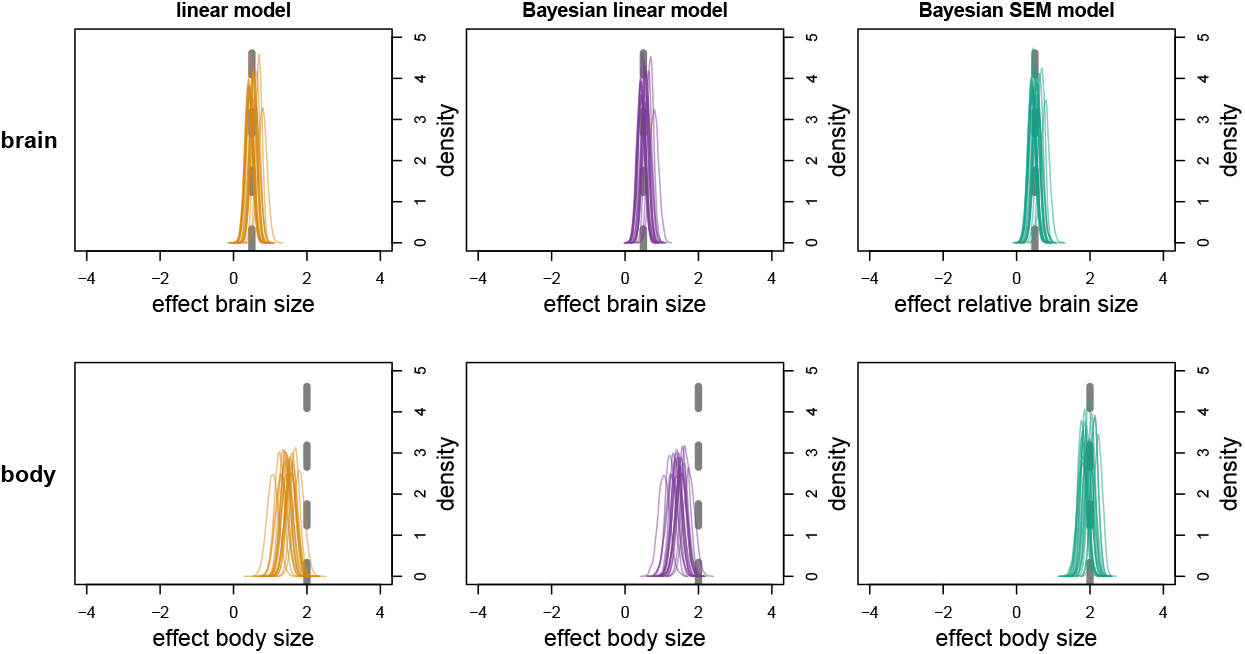
Model with strong effect of body size and weak effect of brain size. Parameter estimates from the linear model, Bayesian linear model and Bayesian strucutral equation model with relative brain size as predictor variable. Dashed grey line is the true value. Orange density plots are normal distributions based on the mean and SE from the linear model. Purple density plots are the posterior distributions from the Bayesian models.

**Figure 13:**
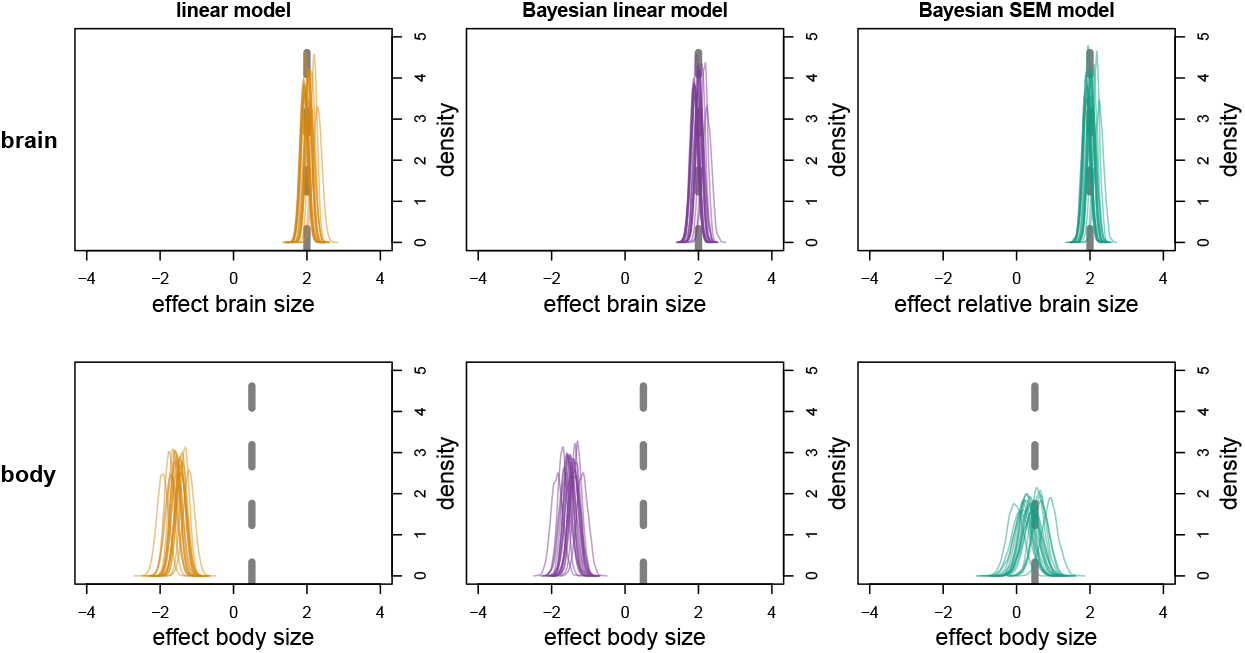
Model with strong effect of brain size and weak effect of body size. Parameter estimates from the linear model, Bayesian linear model and Bayesian strucutral equation model with relative brain size as predictor variable. Dashed grey line is the true value. Orange density plots are normal distributions based on the mean and SE from the linear model. Purple density plots are the posterior distributions from the Bayesian models.

**Figure 14:**
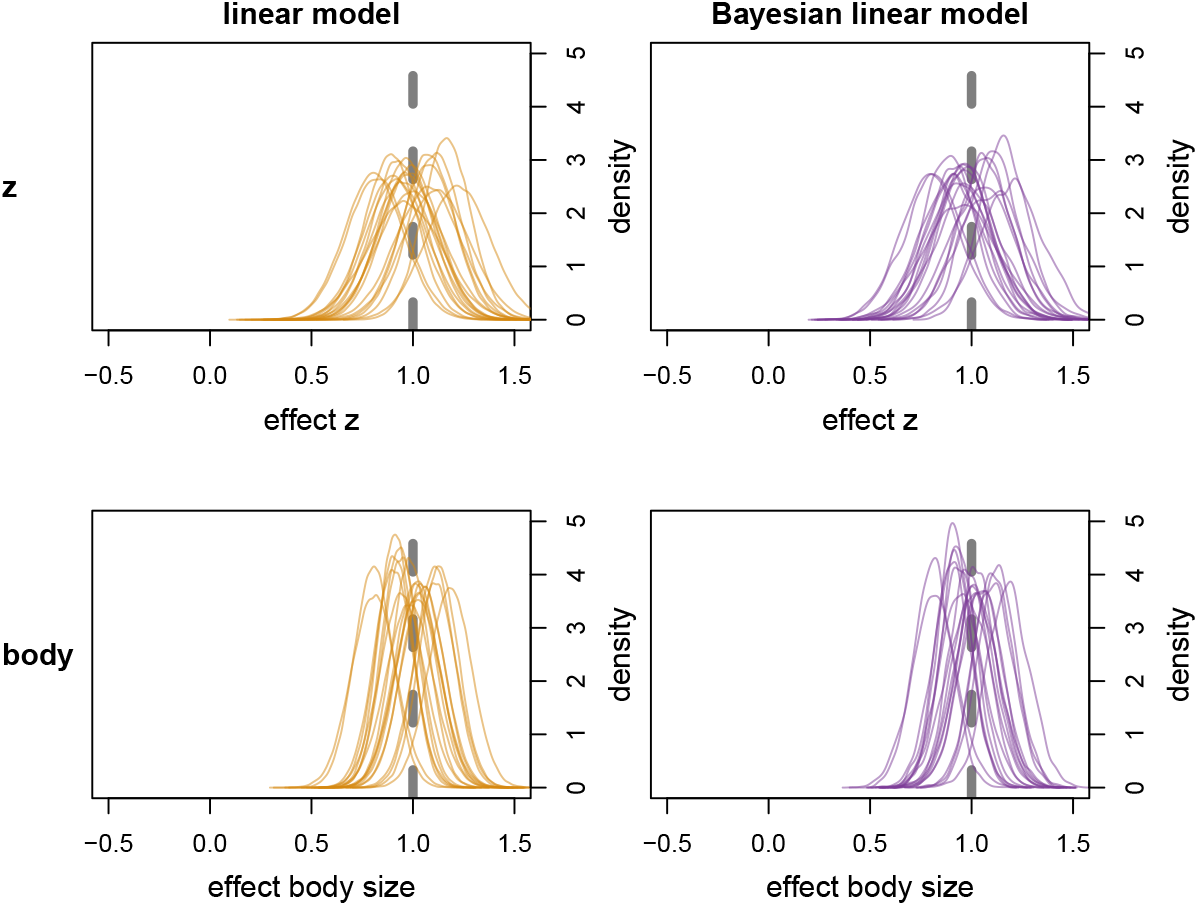
Model with relative brain size as response variable and an additional causal path from body size to z. Parameter estimates from the linear model and Bayesian linear model. Dashed grey line is the true value. Orange density plots are normal distributions based on the mean and SE from the linear model. Purple density plots are the posterior distributions from the Bayesian model.

## Discussion

Several recent studies have claimed to study the effect of relative brain size by including both absolute brain size and body size as predictors (Street et al. 2017; González-Lagos, Sol, and Reader 2010; Isler and Van Schaik 2009; Maklakov et al. 2011; Samia, Pape Møller, and Blumstein 2015; Sol et al. 2022). These studies also reported the direct effect of body size and sometimes drew conclusions based on the sign of this effect. Since all these studies used some version of a linear model (be it phylogenetic and/or Bayesian), they actually tested the effect of absolute brain size, since including body size only accounts for allometry in the response variable. The simulations in this paper showed that including body size as additional predictor to control for allometric scaling of brain size works well if relative brain size is the response variable, but not if relative brain size is a predictor variable. Perhaps counter-intuitively, the effect of brain size was still estimated correctly by all models. It was the body size effect that was biased in the linear models. Simulations with a strong body size effect still found a positive coefficient for this effect, but were biased towards lower values (see Appendix, Figure 12). In simulations with a strong brain size effect the body size coefficients were even negative (see Appendix, Figure 13).

One way to create some intuition about what is going on is that absolute brain size (which was the actual predictor variable included in the linear models) contains information about both body size and relative brain size. In a sense this variable controls for part of the body size effect already. In other words, there are two paths from relative brain size (or the unobserved cognitive ability from the DAG) to *z*: one path is direct and one path is a back-door path through the allometric effect of body size. In another case, where body size itself does not have an effect on *z*, it does not actually need to be included at all (Walmsley and Morrissey 2022). In this simple example, using absolute brain size would be fine. The variation in brain size due to allometric scaling would just create noise. However, in empirical studies the causal structure is often more complex, and researchers should always check if using absolute brain size leads to bias. Using absolute brain size would also lead to a less precise estimate of the effect of brain size, so using relative brain size from a structural equation model would still be preferable.

Structural equation models are similar to phylogenetic path analysis (Gonzalez-Voyer and Hardenberg 2014) and multivariate regressions (Izenman 2013). However, both these techniques do not allow the inclusion of the difference between the observed and predicted brain size from one model as predictor for a second model. A potentially even more powerful approach was put forward by Smaers and Vinicius (2009) and involves reconstructing the ancestral states of both body size and brain size. Recently this approach was used to model brain evolution in mammals and it was shown that relatively large brains can be achieved by divergent paths (Smaers et al. 2021). The authors contest the notion that there is a universal scaling between body and brain size, and instead propose that relatively large brains can just as well be a result of selection towards smaller bodies. Despite this observation, there might still be value in the use of relative brain size for two reasons. First, the largest non-cognitive effects were due to a shift in the slope of the body to brain relation. Such a shift would be less likely to affect results when considering a lower-order taxonomic group (e.g., only primates) and could be partially accounted for by including multiple slopes (e.g., one per family). Second, running a Bayesian multi-regime OU modelling approach is not straightforward and becomes really difficult when many covariates are included, since no step-by-step guide or R package exists. A compromise is to first study the allometric patterns in the taxon of interest and then decide if relative brain size can be used as proxy for cognitive ability.

For this paper I drew all independent variables from normal distributions with zero mean and standard deviation one. I furthermore simulated *z* as a function of relative brain size, rather than simulating both *z* and brain size as a function of the unobserved cognitive ability (as depicted in Figure 1b). I chose to do this to illustrate that even under simple conditions a multiple linear regression cannot estimate the effect of body size correctly. In empirical studies the exact causal structure will be different, but the general observation that one cannot control for allometry in a multiple linear regression when brain size is not the response variable still stands. A regular confounder check as described by Laubach et al. (2021) should inform the design of the model. One example of relative brain size as predictor variable is the study of life history and innovativeness. Sol et al. (2016) suggested that relative brain size predicts both the innovation propensity and maximum life span of birds. They recognised the critique on using residuals, but still used this approach, because they wanted to remove the allometric effects from brain size and not from innovation propensity. This was a valid argument since the original critique assumed a case where *x*_1_ and *x*_2_ are both causing *y* (Freckleton 2002). When *x*_1_ and *x*_2_ are correlated, the use of a residual of *y* ~ *x*_1_ creates a bias. In the case of life history and innovativeness the causal structure is different and this particular problem does not arise. However, in an empirical study measurement error, missing data and phylogenetic covariance still need to be accounted for. With the use of a structural equation model one obtains direct estimates of relative brain size, which includes uncertainty stemming from measurement uncertainty on body size and on brain size. This can also be plotted directly, allowing researchers to visualise how certain the correlation of relative brain size with the third variable is.

Another potential problem with the use of residuals was brought up by Rogell, Dowling, and Husby (2020). They show that many studies have opposite signs between the model with absolute brain size and the model with residual brain size. They argue that coefficients from linear models are difficult to interpret in a biologically meaningful way, when body size and brain size are strongly correlated. In contrasts, Walmsley and Morrissey (2022) show that collinearity does not cause bias per se, and that sign reversal between simple and multiple regressions are simply due to the fact that these answer different biological questions. They do, however, point out that estimating the effect of an agent (*z*) on brain size using multiple regression can be biased if body size is to some extend caused by brain size. As a preliminary solution they propose a Methods of Moments estimator and show that this approach correctly estimates the coefficients in the simulated data. In the current study, the average correlation between body size and brain size was 0.69, which is not as strong as in some empirical studies, but still clearly problematic for the linear models. The SEM estimates did not suffer under these conditions. Adding additional correlation in case I did not bias the parameter estimates of the multiple regressions, but it did increase uncertainty slightly (see Appendix, Figure 14).

Structural equation models allow for the inclusion of measurement error, imputation of missing data and phylogenetic covariance. Smeele et al. (2022) published their R script for such an applied structural equation model, which can be modified to fit most comparative questions. Furthermore, the Bayesian implementation is useful in cases with low sample size. Prior information about parameters can be used to increase accuracy. E.g., it is highly unlikely that an increase in body size would directly lead to a decrease in brain size. The prior for this relationship could therefore be set to only include positive values. This creates a clear advantage over frequentist linear models, which allow any real number and can therefore produce scientifically nonsensical results.

## Conclusion

The aim of this paper was to show that multiple regressions fail to estimate causal effects when relative brain size is a predictor variable. Multiple other approaches can be used instead and the design of the model should be based on the assumed causal structure of the system. I provide a simple structural equation model, which works for the simulated data. Future studies should make sure all back-door paths are closed for their DAG and potentially include additional components to control for measurement error, missing data and phylogeny. The use of these models is not limited to relative brain size, but can be used for any comparative study in which multiple causal paths are of interest.

## Acknowledgements

I would like to thank all anonymous reviewers for valuable input. Their contribution helped to clarify the manuscript and gain new insights.

## Data accessibility

All data is generated in the simulation. Code is publicly available at: https://github.com/simeonqs/Using_relative_brain_size_as_predictor_variable-_serious_pitfalls_and_solutions. Upon acceptance all materials will also be made publicly available on Edmond.

## Appendix

### Supplemental methods

#### Sample size

The main text contained results for data sets with 100 species. To see how the models perform with a smaller or larger dataset, I simulated data for 20 and 1000 species, with 20 simulations per case and sample size.

#### Priors

The main text contained results for Bayesian models with slightly regularising priors. In emperical data sets, one can often choose more informative priors. To test the effect of this I analysed the dataset with 100 species with priors that only allow for positive slopes, with the mean set to the simulated value (1). I also restricted the priors for the intecept parameters to normal(0, 0.25).

To further test the robustness of the models I analysed the data with vague priors, settings intercept and slope to normal(0, 10) and standard variation parameters to exponential(0.1) (note that the mean for the exponential distribution is the inverse of the rate, such that the mean standard deviation here is 10).

#### Effect magnitude

The main text assumes equal magnitude for the body size and brain size effect. To test how the models performed when the body and brain effects on z are unequal in magnitude, I ran two additional simulations with either strong body size effect (*β*_body_ = 2, *β*_brain_ = 0.5) or strong brain size effect (*β*_body_ = 0.5, *β*_brain_ = 2). All other settings and analysis were the same as in the main text.

#### Additional causal path

To test how the multiple regressions can deal with additional causal complexity I simulated a case with brain size as the response variable (similar to case I), but with an additional causal path from body size to *z*. All other settings and analysis were the same as in the main text.

### Supplemental results

